# Dynamic remodeling of mechano-sensing complexes in suspended fibroblast cell-sheets under external mechanical stimulus

**DOI:** 10.1101/2024.11.28.625950

**Authors:** Madoka Suzuki, Keiko Kawauchi, Hiroaki Machiyama, Hiroaki Hirata, Shin’ichi Ishiwata, Hideaki Fujita

## Abstract

Freestanding cell-sheets are valuable bio-materials for use in regenerative medicine and tissue engineering. Because cell-sheets experience various mechanical stimulations during handling, it is important to understand the responses of cells to these stimulations. Here, we demonstrate changes in the localization of various proteins during the stretching of fibroblast cell-sheets. These proteins are known to be involved in mechano-sensing. Upon stretching, actin filaments appear parallel to the stretching direction. At cell-cell junctions, β-catenin forms clusters that co-localize with accumulated vinculin and zyxin as well as the actin filaments. The p130 Crk-associated substrate, known to be present in focal adhesions, is also recruited to these clusters and phosphorylated. Our results suggest that mechano-sensing machinery is formed at cell-cell junctions when the cell-sheets are stretched.

## Introduction

Fabrication of tissue grafts for regenerative medicine relies generally on porous 3D scaffolds made from biocompatible materials such as extracellular matrix (ECM), synthetic polymers, hydrogels or decellularized tissue (Baharvand, Hashemi, Kazemi Ashtiani, & Farrokhi, 2006; Gunatillake & Adhikari, 2003; Ott et al., 2008; Tibbitt & Anseth, 2009). However, these 3D scaffolds often encounter problems, such as insufficient cell density and an immune response to the scaffold materials after implantation. Various scaffold-free tissue engineering technologies have been developed to overcome these issues, such as the use of centrifugal or magnetic force, or 3D printers (Ito et al., 2004; Matai, Kaur, Seyedsalehi, McClinton, & Laurencin, 2020; Mironov, Kasyanov, Markwald, & Prestwich, 2008). Among these, cell-sheet tissue engineering emerged as a promising technology. Freestanding cell-sheets are fabricated using the temperature-sensitive polymer, poly-N-isopropylacrylamide (PNIPAAm). PNIPAAm grafted culture dishes are now becoming popular in the field of regenerative medicine (Okano, Yamada, Sakai, & Sakurai, 1993; Yang et al., 2007). In this technique, enzymatic digestion is not required; instead, cells are detached from the substrate by lowering the temperature below the critical value at which phase transition occurs in PNIPAAm. Therefore, cell-sheets can be obtained while cell-cell interactions remain intact. However, handling of the cell-sheet is difficult owing to the lack of a scaffold, and cell-sheet stretch may occur during tissue fabrication or transplantation, which may induce unwanted mechanical responses to the cells.

Sensing mechanical stimuli is a primary reaction in numerous biological processes, including cell migration, cell differentiation, and cancer metastasis. Ion fluxes through mechano-sensitive channels (Guharay & Sachs, 1984; Sokabe, Sachs, & Jing, 1991), signaling from cell-matrix attachments (Abercrombie & Dunn, 1975; Morgan, Humphries, & Bass, 2007), and signaling from cell-cell junctions (Takeichi, 1995; Yamada, Pokutta, Drees, Weis, & Nelson, 2005) are the major mechano-sensing mechanisms. In mammalian cells, studies into mechano-sensing mostly focus on signaling through focal adhesions (FAs), which consist of protein components including integrins, talin, vinculin, p130 Crk-associated substrate (p130Cas), FA kinase (FAK), and paxillin at the cell-substrate interface (Moore, Roca-Cusachs, & Sheetz, 2010). FAs are attached to stress fibers, which are composed of actin filaments, and so are able to transmit forces generated mainly via myosin II motors on these stress fibers to the underlying substrate.

Cadherins are intercellular adhesion proteins present in the adherens junctions (AJs) between cells. The mechanical forces exerted on AJs are sensed by cadherin-catenin complexes (Borghi et al., 2012; Ladoux et al., 2010; Matsuda et al., 2005) and then transduced into intracellular signaling cascades (Tzima et al., 2005). In epithelial cells, multiple adapter proteins such as α-catenin and vinculin work with cadherin (Ladoux et al., 2010; le Duc et al., 2010; Yonemura, Wada, Watanabe, Nagafuchi, & Shibata, 2010) or through the activation of stretch sensitive Ca^2+^ channels (Ko, Arora, & McCulloch, 2001) to ensure remodeling of the actin cytoskeleton. However, there is still controversy over whether cadherin functions as a mechano-sensor in fibroblasts. One of the major roles of fibroblasts *in vivo* is in ECM production. Mechanical stimulation of fibroblasts results in remodeling of their actin filaments, changes in gene expression, and an increase in the production of ECM (Chiquet, Gelman, Lutz, & Maier, 2009). Increased ECM production enhances matrix rigidity, which subsequently alters gene expression in surrounding cells (Alcaraz et al., 2008; Rizki et al., 2008). This process may regulate cell fate, for example through affecting cell differentiation and tumorigenesis (Butcher, Alliston, & Weaver, 2009).

In this study, the cell responses to mechanical stretching on single-layered cell-sheets fabricated from NIH 3T3 fibroblasts were investigated. Both ends of the cell-sheets were attached to a pair of micro-glass needles and stretched under a fluorescence microscope. We found that stretching induced the formation of actin fibers, which was followed by the relocation of α-catenin, β-catenin, p130Cas and zyxin. This result indicates that cell-sheets rapidly sense mechanical stretching, which may change the cell state when cell-sheets are stretched during handling for tissue fabrication or implantation.

## Results

### Mechanical stretching of cell-sheets

We prepared cell-sheets from NIH3T3 fibroblasts, which express N-cadherin instead of E-cadherin (Reynolds, Daniel, Mo, Wu, & Zhang, 1996). The disruption of cell-sheets in the absence of Ca^2+^ (Supplemental Figure S1A) and immunofluorescence analysis (Supplemental Figure S1B) indicated the involvement of N-cadherins in cell-cell interactions within the cell-sheet. Stress fibers observed on the PNIPAAm-grafted surface (Supplemental Figure S2A) disappeared when cells were detached from the surface by lowering the temperature (Supplemental Figure S2B) and did not reform even when the temperature was returned to 37°C (Supplemental Figure S2C). Instead, actin filaments were localized at the cell-cell boundary regions in the cell-sheets. Actin filaments were observed throughout the cytoplasm after re-attachment to a rigid surface (Supplemental Figure S2D). This observation clearly indicated that actin filament organization is largely influenced by cell attachment to the substrate.

To apply an external force to the floating cell-sheet, we constructed a microscope system composed of two sets of micro-manipulators and a real-time confocal unit serially connected to a dual-view system (Figure 1A, *Materials and Methods*). The cell-sheet attached to the microneedles could be stretched by linearly moving one of the microneedles away from the other (Figure 1B). Cells and cell nuclei deformed in the direction of the stretch and remained deformed until observation was completed (Figure 2, A and B, Supplemental Movie S1). Cells retained their relative positions within the cell-sheet even after stretching (Figure 2A), indicating that cell-cell interactions were maintained throughout the stretch procedure.

**Figure 1.**
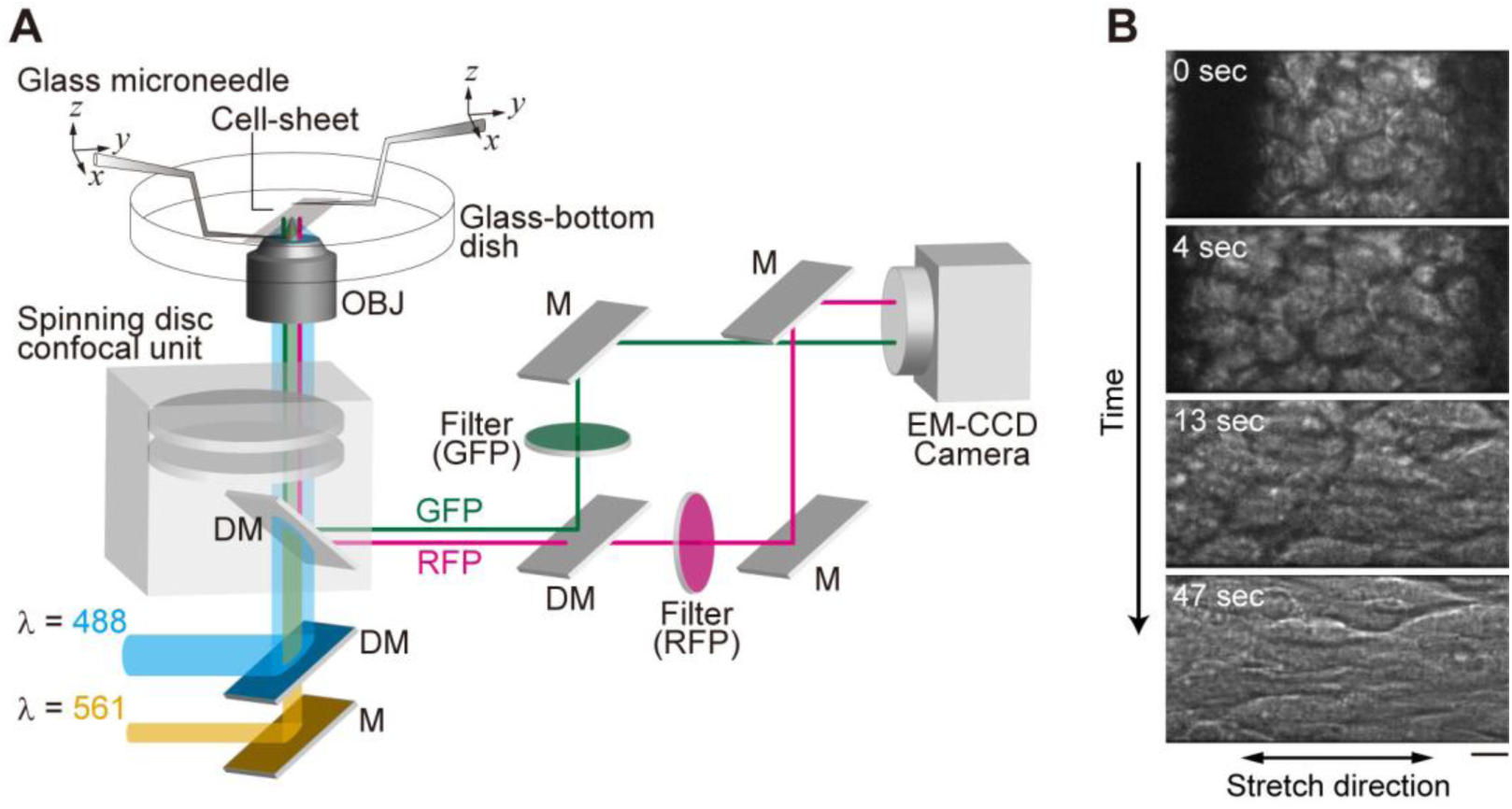
Mechanical stretching of a cell-sheet under the optical microscope. (**A**) Schematic illustration of the optical setup. The cell-sheet was sustained by a pair of glass microneedles attached to both ends of the cell-sheet above a glass-bottom dish. Cells were simultaneously illuminated with blue (488 nm) and yellow-green (561 nm) lasers through a spinning disc confocal unit. GFP and RFP signals were reflected by a dichroic mirror and relayed to a dual-view system to image both signals simultaneously with a single camera. DM, dichroic mirror. OBJ, objective lens. M, mirror. (**B**) Bright-field images of a cell-sheet during stretching. Glass microneedles were located at the left and the right (observed as black shadows in the first and the second images). Double-sided arrow at the bottom indicates the stretching direction. Cell-sheet was stretched to 200% of the slack length. Scale bar, 10 μm.

**Figure 2.**
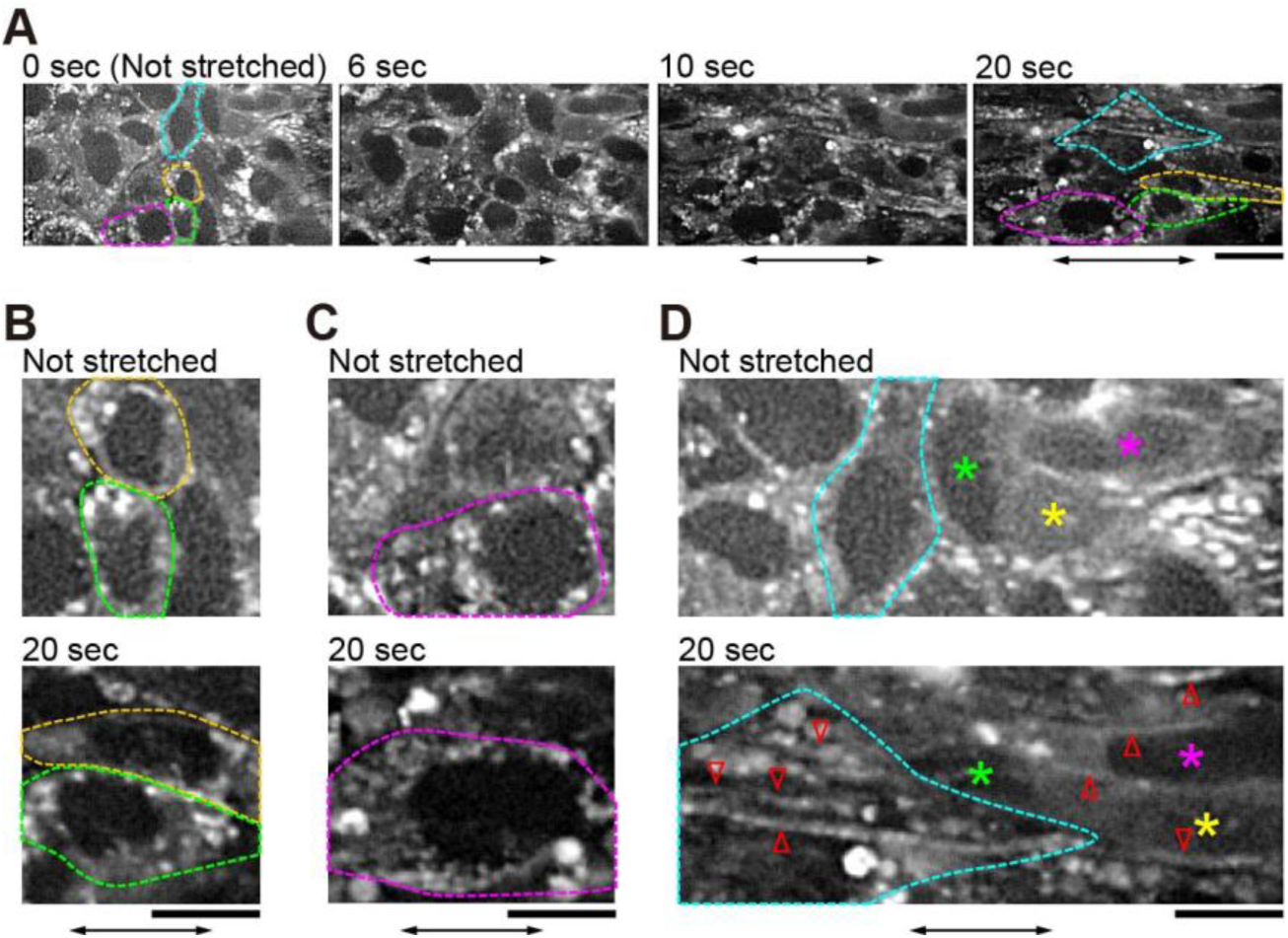
Remodeling of actin filaments observed in a cell-sheet expressing RFP-actin. (**A**) Sequential micrographs of fluorescence images of RFP-actin during stretching. Note that the same cell can continuously be observed. (**B** - **D**) Enlarged images of (***A***) at t = 0 sec (upper panels) and at t = 20 sec (lower panels), illustrating the deformation of cell nucleus (***B***), the increase of dot-like structures after stretching (***C***), and actin filaments newly appeared after stretching (***D***). Cells outlined in the same color or marked by colored stars in (***A*** *- **D***) indicate the same cells. Red arrowheads indicate actin filaments. Double-sided arrows indicate the stretching direction. Scale bars in (**A**) and (**B** - **D**) are 20 and 10 μm, respectively. Independent experiments were performed in 6 cell-sheets from different preparations, all of which gave similar results.

### Actin filament remodeling by mechanical stretch

While the tension on E-cadherin requires actomyosin contraction (Borghi et al., 2012; A. R. Harris et al., 2012), it remains unknown how the forces externally applied at cell-cell junctions affect the remodeling of the actin cytoskeleton. After stretching, the dot-like structure of actin increased in most cells, both in those that previously lacked the structure (Figure 2A, Not stretched) and in those that contained the structure before the stretch (Figure 2C). Actin filaments were clearly visible along the stretch axis (Figure 2D, Supplemental Movie S1, cf. Figure 3A). We confirmed that the newly appeared actin filaments were not those moving in from out-of-focus planes by collecting z-stack images of the cell-sheet before stretching (Supplemental Movies S2 and S3).

**Figure 3.**
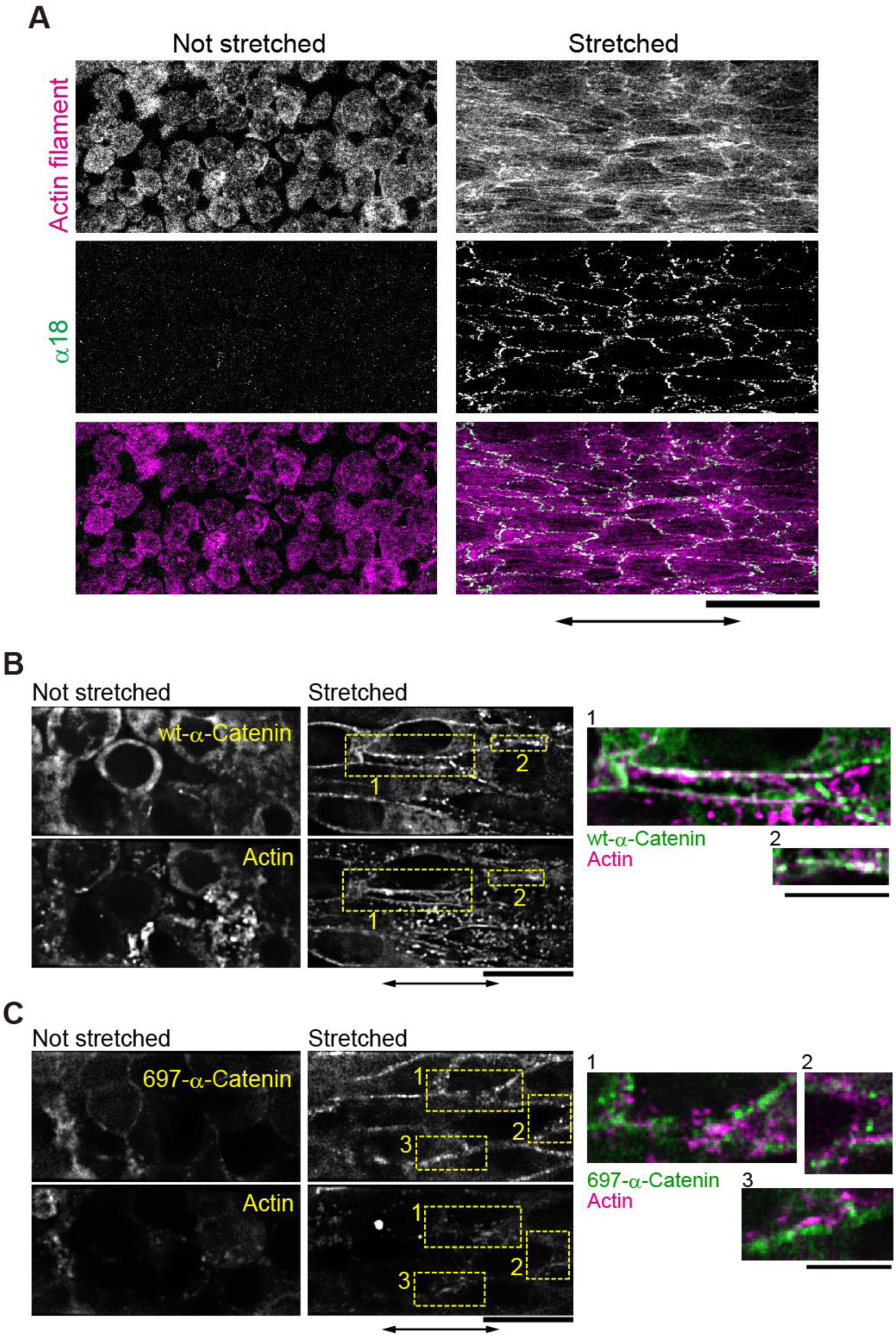
Formation of actin filaments and mechano-sensing by α-catenin in stretched cell-sheet. (**A**) Alexa Fluor 546-conjugated phalloidin was used to visualize actin filaments. Immunofluorescence staining of α-catenin was achieved using a monoclonal antibody, α18, that recognizes α-catenin in a force-dependent manner (see ‘Stretched’). Images show the maximum intensity of the three optical slices from the bottom of the cell sheet. Scale bar, 100 μm. Double-sided arrows indicate the stretching directions. Independent experiments were performed in 2 and 4 cell-sheets from different preparations for non-stretched and stretched conditions, respectively, all of which gave similar results. (**B and C**) Fluorescence micrographs of a cell-sheet expressing GFP-fused wild type α-catenin (GFP-wt-α-catenin) (***B***) or GFP-fused deletion mutant of α-catenin lacking the vinculin binding site (GFP-697-α-catenin) (***C***), and RFP-actin before and after stretching, held for 30 min. The overexpression of wt-α-catenin did not interfere with the formation of actin filaments upon stretching (***C***). Right panels are the magnified and merged views of the areas indicated by yellow rectangles to the left. Scale bars, 20 and 10 μm in left and right panels, respectively. Independent experiments were performed in 8 cell-sheets in (***B***) and 4 cell-sheets in (***C***) from different preparations, all of which gave respectively similar results.

### Accumulation of AJ proteins at cell-cell junctions by mechanical stretch

Next, we investigated how tension was transmitted into the cells via cell-cell junctions in the cell-sheet. In AJs, vinculin binds to actin filaments and α-catenin (Huveneers & de Rooij, 2013). The vinculin-binding site in α-catenin is considered to open upon stretching (Yonemura et al., 2010). Consistent with these reports, the antibody α18, which recognizes α-catenin under tension (Nagafuchi & Tsukita, 1994), bound to cell-cell junction sites in the cell-sheet upon stretching (Figure 3A). Furthermore, introduction of a deletion mutant of α-catenin lacking the vinculin binding site (697-α-catenin) into cells impaired the formation of actin filaments upon stretching (Figure 3, B and C, Supplemental Movies S4 and S5). Tension also changed the distribution of AJ-associating proteins, such as β-catenin and vinculin. Beta-catenin, which was distributed homogeneously at the cell-cell junctions under the non-stretched condition, formed clusters soon after stretching was applied (Figure 4A). In areas where β-catenin clusters appeared, accumulation of actin filaments (Figure 4, A and B, Supplemental Movie S6) and vinculin was observed (Figure 4, D and E, Supplemental Movies S7 and S8). These clusters resemble those observed previously in fibroblasts and epithelial cells, which were composed of P-cadherin, α/β-catenin, ZO-1 and actin (Adams, Nelson, & Smith, 1996; Vasioukhin, Bauer, Yin, & Fuchs, 2000; Yonemura, Itoh, Nagafuchi, & Tsukita, 1995). As punctate AJs were formed in cells on a rigid surface (Supplemental Figure S3, A-E), the clusters of α/β-catenin and vinculin formed in stretched cell-sheets often appeared in a periodic manner (Supplemental Figure S3, A-E).

**Figure 4.**
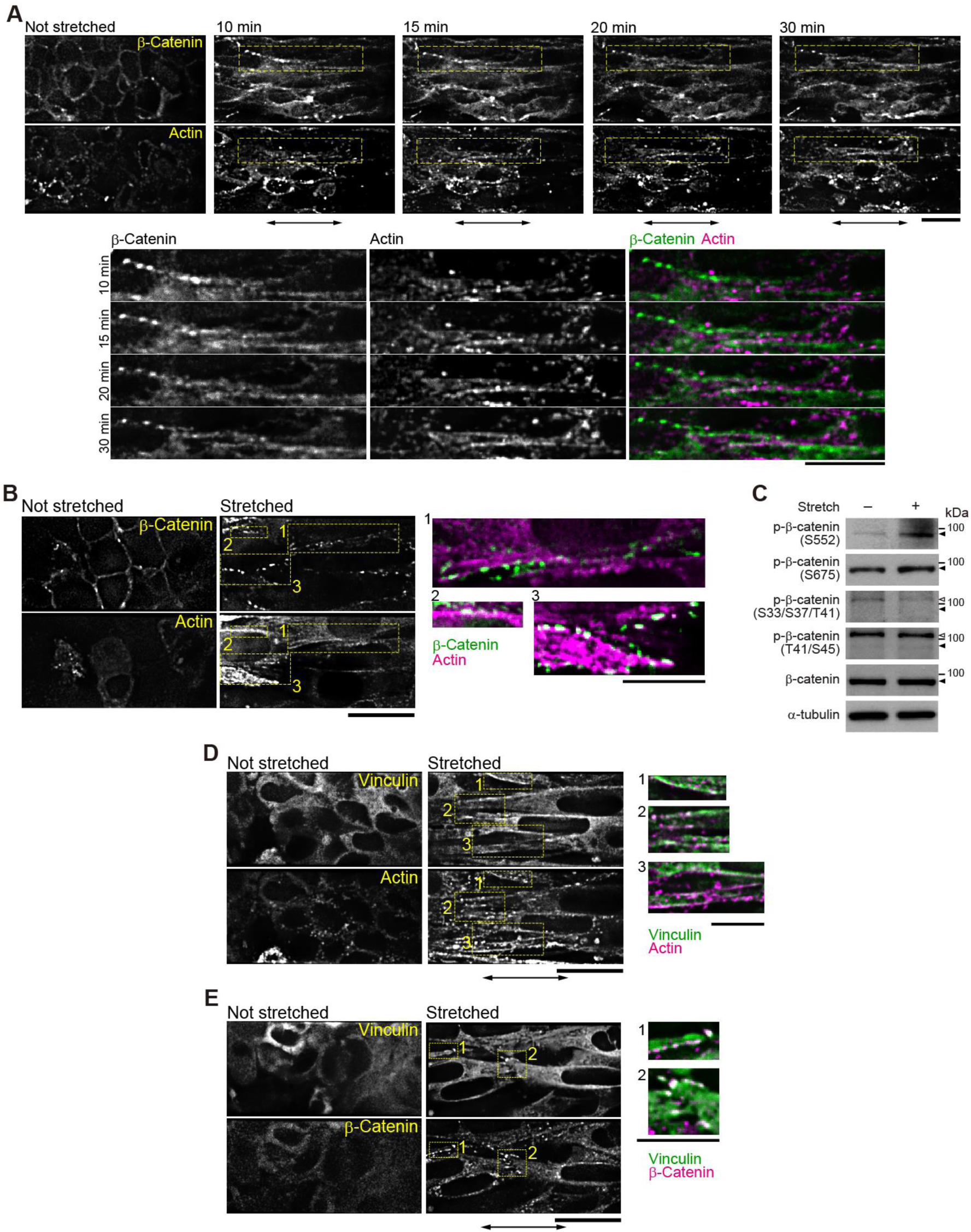
Assembly of adherens junction proteins observed in cell-sheets. Two fluorescent proteins were simultaneously observed on a single camera. Double-sided arrows indicate the stretching direction. (**A**) (Top) Sequential micrographs of a cell-sheet expressing GFP-β-catenin and RFP-actin before and after stretching, held for 10, 15, 20 and 30 min. (Bottom) Magnified views of the areas indicated by yellow rectangles in the top panels. Scale bars, 20 μm. (**B**) Fluorescence micrographs of a cell-sheet expressing GFP-β-catenin and RFP-actin before and after stretching, held for 30 min. For better S/N ratio, image was obtained only once before stretching and 30 min after stretching. The 17 cell-sheets out of 19 independent experiments from different preparations gave similar results. (**C**) Western blot analysis using antibodies against phospho-β-catenin (S552), phospho-β-catenin (S675), phospho-β-catenin (S33/S37/T41), phospho-β-catenin (T41/S45), β-catenin and α-tubulin as a loading control. Extracts were obtained from cell-sheets before and after induction of stretching and held for one minute. Filled and open arrow heads indicate β-catenin and nonspecific bands, respectively. (**D** and **E**) Fluorescence micrographs of a cell-sheet expressing GFP-vinculin and RFP-actin (***D***) or RFP-β-catenin (***E***) before and after stretching, held for 30 min. The 5 cell-sheets out of 8 and 6 cell-sheets out of 9 independent experiments from different preparations gave similar results in (***D***) and (***E***), respectively. In (**B, D** and **E**), right panels are magnified and merged views of the areas indicated by yellow rectangles to the left. Scale bars, 20 and 10 μm in left and right panels, respectively.

Moreover, stretching influenced the phosphorylation status of β-catenin. N-cadherin as well as E-cadherin interactions mediate the transduction of mechanical stress into cellular signaling pathways (Ganz et al., 2006). Beta-catenin, which directly binds to both the intracellular domain of cadherins and the actin-binding protein α-catenin, is a mediator of mechano-transduction at cadherin-based adhesions (Desprat, Supatto, Pouille, Beaurepaire, & Farge, 2008; Morales-Camilo et al., 2024; Roper et al., 2018). It is known that phosphorylation of β-catenin at Ser 552 (Fang et al., 2007) or Ser 675 (Hino, Tanji, Nakayama, & Kikuchi, 2005; Taurin, Sandbo, Qin, Browning, & Dulin, 2006) enhances its signaling activity and causes the localization of β-catenin to cell-cell contacts (Maher, Mo, Flozak, Peled, & Gottardi, 2010), whereas β-catenin phosphorylated at S33/ S37/T41 or T41/S45 cannot associate with cadherins (Maher et al., 2010). In the fibroblast cell-sheet, phosphorylation at both S552 and S675 of β-catenin increased within one minute of stretching, while phosphorylation at S33/S37/T41 and T41/S45 was unaffected (Figure 4C).

Our results suggest that mechanical stress in the cell-sheet stretches α-catenin, exposes the vinculin binding site and enhances the attachment of actin filaments to the cell-cell junction complex. The external tension also stimulates the phosphorylation of β-catenin at S552 and S675, resulting in signal transduction from cell-cell junctions to intracellular pathways.

### Accumulation of FA proteins at cell-cell junctions by mechanical stretch

FA proteins such as FAK and paxillin are recruited to cell-cell junctions upon mechanical stimulation, and p130Cas co-localizes with nephrocystin at cell-cell junctions in epithelial cells (Birukova, Malyukova, Poroyko, & Birukov, 2007; Donaldson et al., 2000; Sun et al., 2009). Given that the formation of actin filaments and remodeling of AJ proteins were induced upon mechanical stretch, FA proteins in the fibroblast cell-sheet may also be recruited to cell-cell junctions in the absence of FAs and interact with AJ proteins in response to stretching. In line with this hypothesis, p130Cas formed clusters at cell-cell junctions along actin filaments and co-localized with β-catenin clusters (Figure 5, A and B, Supplemental Movies S9 and S10). Such the localization is in contrast to that of p130Cas at FA on a rigid surface (Supplemental Figure S3, F and G). Furthermore, the phosphorylation of both p130Cas and FAK increased within one minute of stretching (Figure 5C). These results suggest that p130Cas and FAK are recruited to the sites of force-bearing cell-cell junctions.

**Figure 5.**
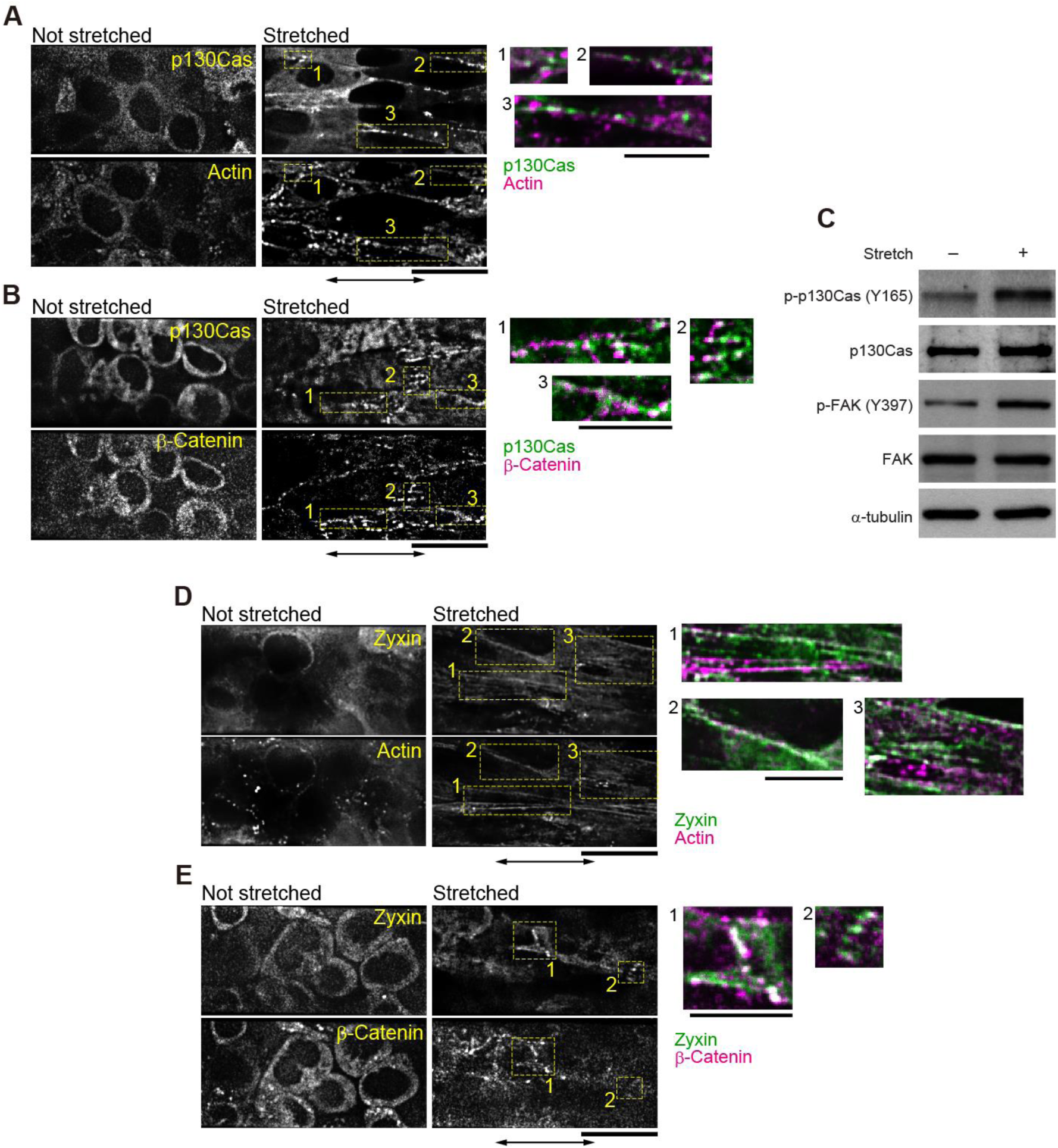
Assembly of focal adhesion proteins observed in cell-sheets. (**A** and **B**) Fluorescence micrographs of a cell-sheet expressing GFP-p130Cas and RFP-actin (***A***) or mCherry-β-catenin (***B***) before and after stretching, held for 30 min. The 8 cell-sheets out of 10 and 10 cell-sheets out of 12 independent experiments from different preparations gave similar results in (***A***) and (***B***), respectively. (**C**) Western blot analysis using antibodies against phospho-p130Cas (Y165), p130Cas, phospho-FAK (Y397), FAK and α-tubulin as a loading control. Extracts were obtained from cell-sheets before and after stretching and held for one minute. (**D** and **E**) Fluorescence micrographs of a cell-sheet expressing GFP-zyxin and RFP-actin (**D**) or mCherry-β-catenin (**E**) before and after stretching, held for 30 min. Independent experiments were performed in 9 cell-sheets in (***D***) and 6 cell-sheets in (***E***) from different preparations, all of which gave respectively similar results. In (**A, B, D** and **E**), the right panels are the magnified and merged views of the areas indicated by yellow rectangles to the left. Scale bars, 20 and 10 μm in left and right panels, respectively.

Stretching also affected zyxin localization. Zyxin is a protein present in both FAs (Supplemental Figure S3H) and AJs (Supplemental Figure S3I). The localization of zyxin to FAs depends on the mechanical force in fibroblasts and epithelial cells (Colombelli et al., 2009; Hirata, Tatsumi, & Sokabe, 2008; Nguyen, Uemura, Shih, & Yamada, 2010), and its presence at cadherin adhesion sites with actin, α/β-catenin and vinculin has also been observed in epithelial cells (Hansen & Beckerle, 2006; Nguyen et al., 2010; Vasioukhin et al., 2000). Here, when the fibroblast cell-sheet was stretched, zyxin localization changed from a broad distribution within the cytosol to clustered structures (Figure 5D, Supplemental Movie S11). In the same region, actin filaments extended in the direction of the stretch and β-catenin clusters were also present (Figure 5E, Supplemental Movie S12). Zyxin is known to promote E-cadherin-mediated actin assembly at AJs (Nguyen et al., 2010), suggesting that the recruitment of zyxin by externally applied force promoted actin polymerization at N-cadherin-mediated cell-cell junctions in the fibroblast cell-sheet.

## Discussion

Actomyosin contractility is required for mechano-transduction through E-cadherin/catenin complexes in epithelial cell-sheets (A. R. Harris et al., 2012). Since stress fibers are more abundant in fibroblasts than in epithelial cells on a rigid surface, force-dependent formation and remodeling of the actin cytoskeleton may be more sensitive in fibroblasts. Here, to the best of our knowledge, the effect of mechanical stretching on suspended fibroblast cell-sheets was reported for the first time. We demonstrated that stress fibers disappeared when the fibroblast cell-sheets detached from the substrate (Figure 2, Supplemental Movie S2). Integrin adhesion receptors recruit FA proteins, including p130Cas and FAK, upon binding to the extracellular matrix. This process further activates Rho GTPases, which induces actin polymerization and actomyosin contraction, and the actin cytoskeleton also regulates integrin activation (Pelham & Wang, 1997; Shemesh, Geiger, Bershadsky, & Kozlov, 2005; Wehrle-Haller & Imhof, 2002). The loss of FAs upon detachment from the substrate is therefore very likely to impair stress fiber formation, as we observed here.

When the cells were stretched and maintained, actin filaments were observed (Figs. 2 - 5). These actin structures resemble those found in endothelial cells in tissues exposed to high mechanical forces (White, Gimbrone, & Fujiwara, 1983; Wong, Pollard, & Herman, 1983). Dot-like and filamentous structures appeared even during stretching (Figure 2A, Supplemental Movie S1). It is therefore probable that these structures were initially formed by the rearrangement of existing actin filaments (Hirata, Tatsumi, & Sokabe, 2007) and then by the polymerization of actin monomers.

We observed that β-catenin (Figure 4, A, B and E, Supplemental Movie S6) and vinculin (Figure 4, D and E, Supplemental Movie S7) clusters appeared after the stretch, and that each cluster of β-catenin and vinculin co-localized with actin filaments (Figure 3, B and D, Supplemental Movies S6 and S7) and with each other (Figure 4E, Supplemental Movie S8). Therefore, these three proteins likely co-localize when mechanical force is present. Beta-catenin and vinculin are both components of the AJs. AJs are formed by cadherins connected to the cytosolic actin filament network (Abe & Takeichi, 2008; T. J. Harris & Tepass, 2010; Taguchi, Ishiuchi, & Takeichi, 2011). Previous reports have shown that vinculin senses mechanical stimuli and is recruited to cadherin-positive structures at cell-cell junctions under force (Huveneers & de Rooij, 2013; Twiss et al., 2012; Yonemura et al., 2010). Considering the co-localization of proteins that are known to bind to each other, it is likely that these proteins form a complex resembling that of an AJ in the fibroblast cell-sheet upon stretching.

The substrate domain of p130Cas, which is characterized by fifteen YXXP motifs, is phosphorylated by Src at the FA sites (Moore et al., 2010). Mechanical forces expose the YXXP motif of p130Cas and enhance its susceptibility to phosphorylation (Sawada et al., 2006). Here, we found that in fibroblast cell-sheets p130Cas did not localize to any specific area under no-force conditions (Figure 5, A and B). However, upon stretching, p130Cas co-localized with actin and β-catenin (Figure 5, A and B, Supplemental Movies S9 and S10). Taken together, it is plausible that the phosphorylation level of p130Cas increased (Figure 5C), following which actin assembly was induced at the sites of force-bearing cell-cell junctions.

In this study, we showed that clustered structures composed of β-catenin, vinculin, zyxin, p130Cas and actin filaments appear at cell-cell junctions upon the application of stretching. Similar structures composed of cadherin, α-catenin, vinculin and actin filaments have been observed at cell-cell adhesion sites in epithelial and endothelial cells (Huveneers et al., 2012; Millan et al., 2010; Yonemura et al., 1995). Punctate structures composed of E-cadherin and actin filaments have also been reported in epithelial cells (Adams, Chen, Smith, & Nelson, 1998). Although we did not image cadherins in our study and earlier studies did not identify p130Cas at these sites, similarities in appearance and protein composition suggest that these structures are equivalent. Notably, previous studies have shown that both FA and AJ components including zyxin, β-catenin, talin, paxillin and cadherins are recruited to highly tensed actin filaments both in vitro and in living fibroblasts (Kiyoshima, Tatsumi, Hirata, & Sokabe, 2019; Smith et al., 2010; Sun et al., 2020). Thus, it is probable that intricate structures composed of cadherin, α/β-catenin, vinculin, zyxin, p130Cas and actin filaments are formed at cell-cell junctions in the fibroblast cell-sheet in the presence of an external force.

The response to mechanical stimului is frequently studied in cells cultured on an elastic membrane, or by applying shear stress using laminar flow. Our method can serve as a 2D pseudo-tissue model for studying the effects of mechano-stimuli during the handling of fabricated tissue grafts, thereby providing insights into cell state changes during tissue fabrication procedures or implantation.

## Materials and Methods

### Cell culture

NIH 3T3 cells were purchased from Riken Cell Bank (Riken, Tsukuba, Ibaraki, Japan) and cultured in DMEM (11965, Thermo Fisher Scientific, Waltham, MA, USA) supplemented with 10% fetal bovine serum (FBS; 10279, Thermo Fisher Scientific), penicillin-streptomycin (100 units ml^-1^, 15140, Thermo Fisher Scientific) at 37°C with 5% CO_2_. Cells were routinely checked for mycoplasma contamination using MycoAlert detection kit (Lonza, Walkersville, MD, USA). For preparation of NIH3T3 cells expressing fluorescent proteins, retroviral vectors were prepared as follows. Using a mouse spleen cDNA library (9536, TaKaRa, Kusatsu, Shiga, Japan), cDNAs of β-catenin, vinculin, actin, α-catenin and α-catenin lacking vinculin binding site (α-catenin697) were amplified by PCR and cloned into pcDNA3 vector (V79020, Thermo Fisher Scientific) with a cDNA of GFP or RFP, then subcloned into the retroviral vector pBabe. GFP-p130Cas pBabe was a kind gift from Dr. Yasuhiro Sawada. The cloned constructs were cotransfected with the packaging plasmids to HEK293T cells by FuGene6 according to manufacturer’s instruction. Retroviral supernatants were harvested 2 days after transfection and applied to NIH3T3 cells. Infected cells were selected using puromycin (100 ng ml^-1^, P-1033, AG Scientific, San Diego, CA, USA) and hygromycin B (200 μg ml^-1^, 10687, Thermo Fisher Scientific) for 2-3 days.

### Cell-sheet preparation

Cells were seeded onto UpCell dishes two days before carrying out the stretch experiment to allow them to reach 100% confluence. For observation under an optical microscope, cells were seeded onto 35 mm dishes (CS3007, CellSeed, Tokyo, Japan). For immunoblot analysis, cells were seeded onto 60 mm dishes (CS3006, CellSeed). Cell-sheets were prepared by first removing the cell culture media, then washing with 1 × PBS, pH 7.4, and finally adding 3 ml of Leibovitz’s CO_2_-independent media (21083, Thermo Fisher Scientific) without phenol red supplemented with 10% FBS. The whole process was performed on a hot plate (KM-1, Kitazato Corporation, Shizuoka, Japan) set to 40°C. The temperature was then decreased to 28°C to allow detachment of the cell-sheet from the UpCell dish. Full detachment of the cell-sheet from the UpCell dish takes approximately 20 to 30 minutes at 28°C.

### Needle preparation

Glass rods (G-1000, Narishige, Tokyo, Japan) were used to prepare the needles for stretching of the cell-sheets. Heat was applied to the middle of the glass rod before quickly pulling the heated rod swiftly at both ends to obtain the desired diameter at its center. Pulled glass rods were left to cool, cut into half and trimmed to the desired length. Each needle was prepared by heating and bending a glass rod at two points (as illustrated in Figure 1*A*), producing a 120° angle at point 1 and an angle of slightly less than 90° at point 2. Needles were coated using Cell-Tak Cell and Tissue Adhesive (354241, BD Biosciences, Franklin Lakes, NJ, USA), cell adhesion supporting protein purified from *Mytilusedulis*. Cell-Tak solution was prepared using 0.1 M Sodium Bicarbonate (37116-00, Kanto Chemical, Tokyo, Japan), pH 8.0, and 1 N Sodium Hydroxide (S2770, Sigma-Aldrich, St. Louis, MO, USA), using the recommended ratio of 10 μl of Cell-Tak, 285 μl of Sodium Bicarbonate, pH 8.0, and 5 μl of 1 N NaOH (added immediately before coating) to make 300 μl of Cell-Tak solution. Three ml of Cell-Tak solution was prepared for 10 to 12 needles. Needles were incubated in the Cell-Tak solution for 30 min at room temperature. Needles were then washed with distilled water to remove Sodium Bicarbonate, air-dried and stored at 4°C.

### Optical setup of dual-color real-time confocal microscopy

Optical setup (Figure 1*A*) was built around the inverted microscope (IX81, Olympus, Tokyo, Japan). For the purpose of simultaneous imaging of two fluorescent proteins, a 488 nm laser (CUBE 488, Coherent, Santa Clara, CA, USA) and a 561nm laser (Sapphire 561, Coherent) were guided into a spinning disc confocal unit (CSU-10, Yokogawa, Tokyo, Japan) attached to the left side port of the microscope. Cells were observed using an oil immersion objective (APON 60XOTIRF, Olympus). The fluorescence images of GFP, and RFP or mCherry were separated using a dichroic mirror (FF580-FDi01, Semrock, Rochester, NY, USA) placed after the confocal unit, passed through respective band pass filters, FF01-520/35 (Semrock) for GFP and FF01-607/36 (Semrock) for RFP or mCherry, and projected onto a single EM-CCD camera (iXonEM+ DU-897, Andor Technology, Antrium, UK). Optical slices were captured along the *z*-axis every 0.198 μm using the Andor iQ software (Andor Technology). Deconvolution of confocal fluorescence micrographs in Figs. 2-5 and Supplemental Movies S1-S12 were processed by AutoQuant software (Roper Industries, Sarasota, FL, USA).

### Stretching of cell-sheets under the optical microscope

Detached cell-sheets were transferred to 60 mm glass-bottom dishes for live-cell imaging under the optical microscope. Cell-sheets were cultured in Leibovitz’s CO_2_-independent media (21083, Thermo Fisher Scientific) without phenol red, supplemented with 10% FBS, during live-cell imaging. Needles (ϕ = 100~140 μm) were positioned using a motorized micromanipulator (EMM-3NV, Narishige). The two needles were adjusted parallel to each other and the distance between them set at 300-400 μm. Needles were lowered until their tips were attached to the cell-sheet. After 5-10 minutes the cell-sheet bound to the needles was lifted up so that the cells were no longer in contact with the bottom of the dish. The temperature of the culture dish was adjusted using the stage top incubator and the heater for the objective lens (INUB-ONICS, Tokai Hit, Shizuoka, Japan). Cell-sheet stretching was performed after the initial z-scan and the stretching was carried out within one minute. In the majority of our experiments, cell-sheets were stretched to 200% of that before the stretch. To reduce the effect of the cell-needle attachments, observations were made at a substantial distance away from the microneedles (more than 100 μm).

### Immunoblot analysis

Needles (ϕ = 0.5~1 mm) were positioned at both ends of a cell-sheet. The cell-sheet was then stretched and held for one minute. The stretched cell-sheet, still attached to both needles, was then washed with cold 1× PBS followed by the addition of 500 μl of a lysis buffer [50 mM Tris, pH 8.0, 150 mM NaCl, 1% Triton-X100, 0.5% SDS, 10 mM EDTA, 1 mM Na_3_VO_4_, 10 mM NaF, protease inhibitor cocktail (04693159001, Roche, Basel, Switzerland) and 1 mM DTT]. Needles were removed and the lysate was collected, sonicated and centrifuged at 20,000 ×*g* for 20 minutes. The supernatants were then subjected to SDS–PAGE. Proteins were transferred to PVDF membrane and probed with the indicated antibodies at 1:1000 dilution. Anti-phospho-β-catenin S552 (#9566), anti-phospho-β-catenin S675 (#4176), anti-phospho-β-catenin S33/S37/T41 (#9561), anti-phospho-β-catenin T41/S45 (#9565), anti-β-catenin (#9562), anti-phospho-p130Cas Y165 (#4015), anti-FAK (#3285) (Cell Signaling Technology, Danvers, MA, USA), anti-p130Cas (clone 2C1.1; Merck, Darmstadt, Germany), anti-phospho-FAK Y397 (#07-829) (Merck), and anti-α-tubulin (clone B-5-1-2;Sigma-Aldrich) antibodies were used for immunoblotting.

### Immunofluorescence microscopy

For immunofluorescence staining of α-catenin by α18 that recognizes α-catenin under tension (Nagafuchi & Tsukita, 1994), cells were fixed with 1% PFA, permeabilized with 0.2% Triton X-100, and then blocked with 5% BSA in PBS. Alexa Fluor 488-conjugated goat anti-rat IgG (A11006, Thermo Fisher Scientific) was used as a secondary antibody. Alexa Fluor 546-conjugated phalloidin (A22283, Thermo Fisher Scientific) was used to stain F-actin. Images were acquired using a 60× objective (PLAPON 60XO, Olympus) and a laser scanning confocal microscope (FluoView FV1000, Olympus).

## Abbreviations used

AJ: adherens junction
ECM: extracellular matrix
FA: focal adhesion
FAK: focal adhesion kinase
p130Cas: p130 Crk-associated substrate
PNIPAAm: poly-N-isopropylacrylamide

## Acknowledgements

The authors thank Prof. M.P. Sheetz (National University of Singapore) for fruitful discussion. This work was supported by PRESTO, the Japan Science and Technology Agency Grant Number JPMJPR15F5 (to M.S.) and by the Japan Society for the Promotion of Science (JSPS) KAKENHI Grant Numbers 23687021 (to M.S.), 22227005 (to S.I.), and 24570191 (to H.F.).

