## Supplementary material for "Dynamic remodeling of mechano-sensing complexes in suspended fibroblast cell-sheets under external mechanical stimulus": Suppliment info

#### Supplemental Figures

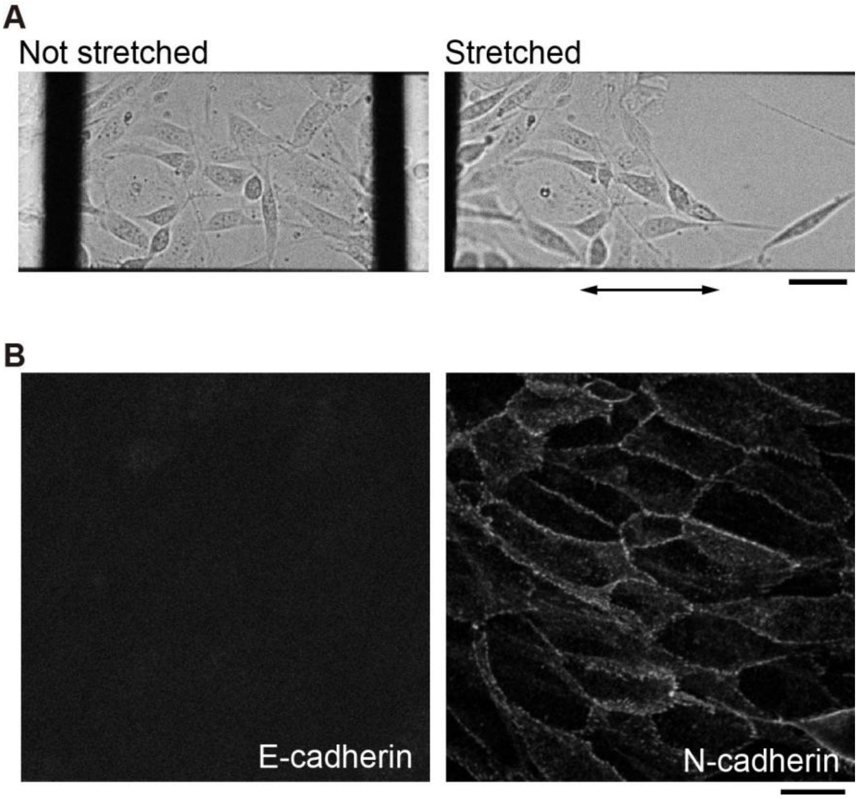


**Supplemental Figure S1.**

(**A**) Cell-cell junctions in the cell-sheet are loose (left panel) and easily broken by mechanical stretch (right panel) in the presence of EGTA. Glass needles were attached to the cell sheet 20 min after addition of 4 mM EGTA. Double-sided arrow indicates the direction of the stretch. Scale bar, 40 $\mu$m. (**B**) NIH3T3 cells stained against E-cadherin (left panel) and N-cadherin (right panel). NIH3T3 cells were cultured until confluence on a glass-bottom dish. Scale bar, 20 $\mu$m.

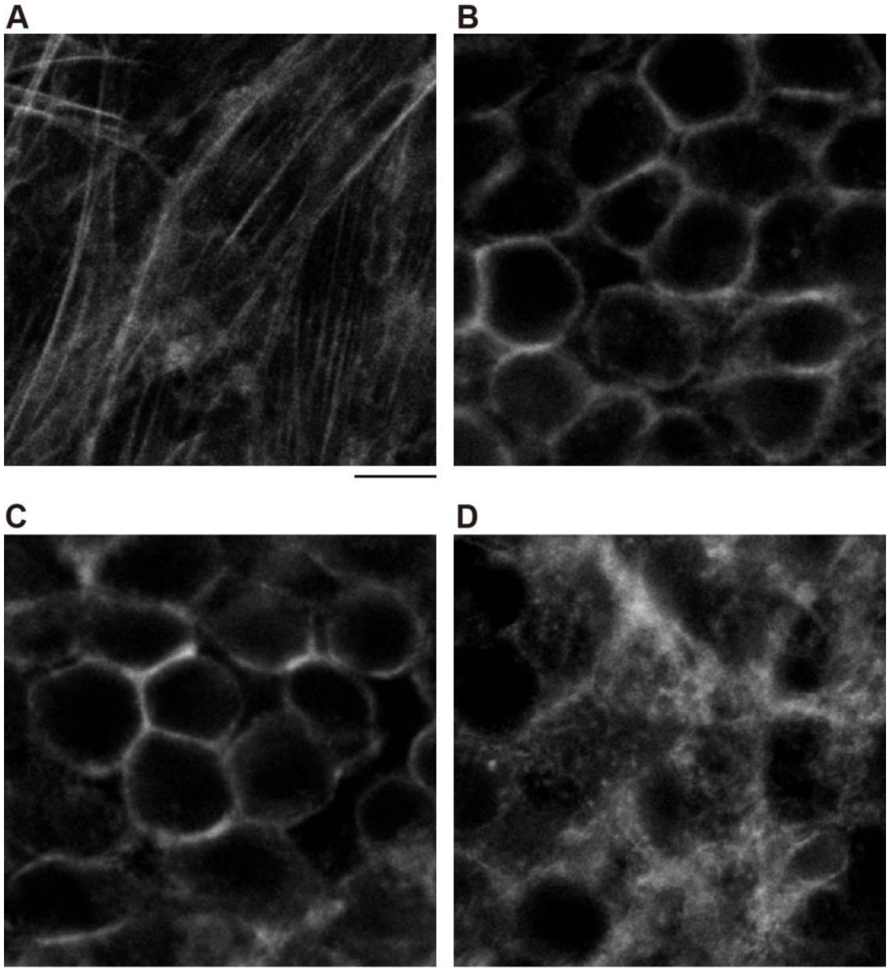


**Supplemental Figure S2.**

NIH3T3 cells stained against actin filaments using Alexa568-phalloidin after 4% paraformaldehyde fixation. (**A**) Cells cultured on a rigid surface. Cells were cultured until confluence. (**B**) Cell-sheet just after fabrication. (**C**) Cell-sheet fabricated and cured for 30 min at 37°C. (**D**) Cell-sheet re-attached to a rigid surface for 5 min. Scale bar, 20 μm.

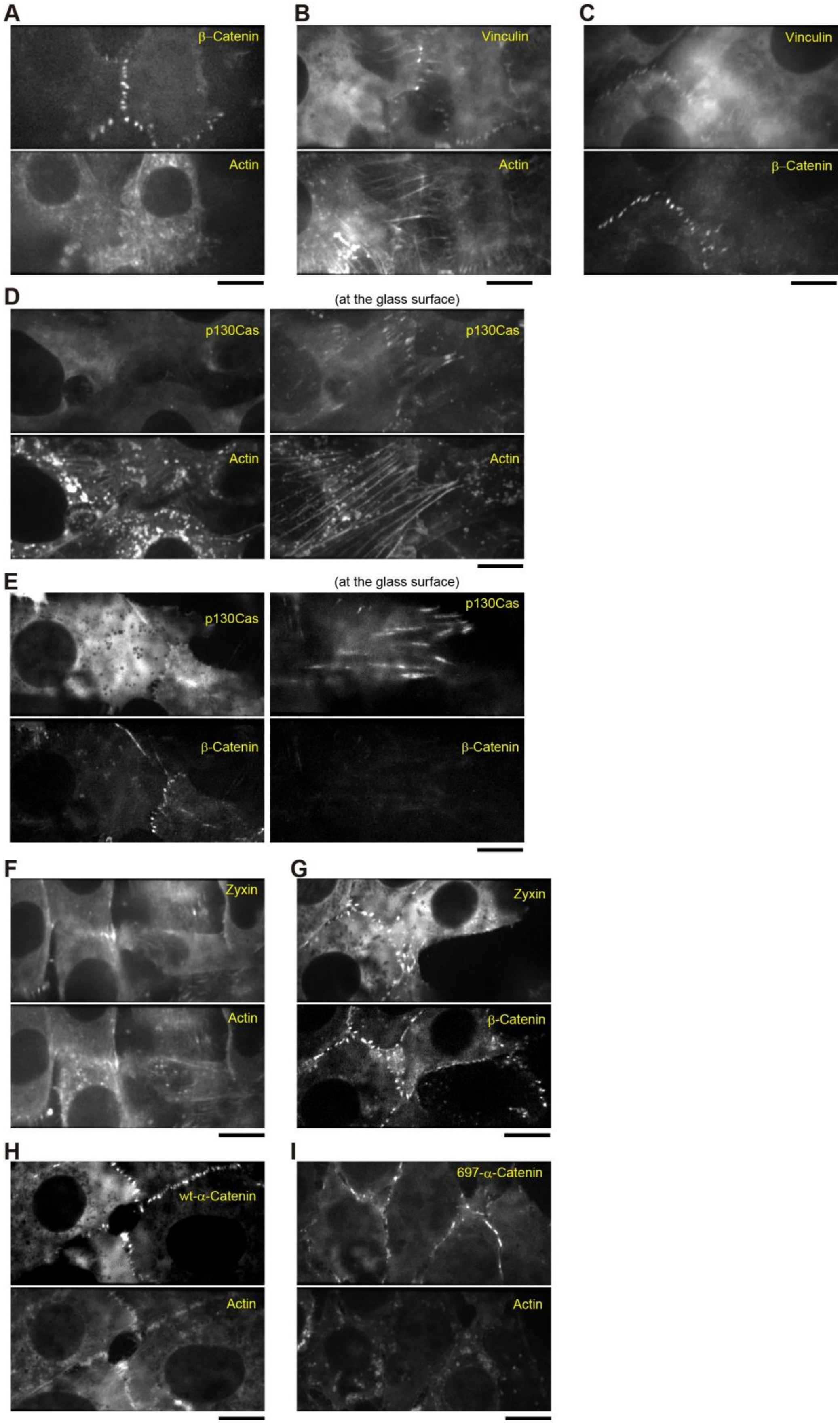
A
β–Catenin
Actin
B
Vinculin
Actin
C
Vinculin
β–Catenin
D
(at the glass surface)
p130Cas
p130Cas
Actin
Actin
E
(at the glass surface)
p130Cas
p130Cas
β-Catenin
β-Catenin
F
Zyxin
Actin
G
Zyxin
β-Catenin
H
wt-α-Catenin
Actin
I
697-α-Catenin
Actin

28

#### Supplemental Figure S3.

Assembly of adherens junction and focal adhesion proteins observed in cells cultured on a rigid glass surface. Two fluorescent proteins were simultaneously observed on a single camera. Scale bars, 20 μm. Fluorescence micrographs show cells expressing GFP-β-catenin and RFP-actin (**A**), GFP-vinculin and RFP-actin (**B**), GFP-vinculin and mCherry-β-catenin (**C**), GFP-p130Cas and RFP-actin (**D**), GFP-p130Cas and mCherry-β-catenin (**E**), GFP-zyxin and RFP-actin (**F**), GFP-zyxin and mCherry-β-catenin (**G**), GFP-wt-α-catenin and RFP-actin (**H**), and GFP-697-α-catenin and RFP-actin (**I**). Images in (**A** - **C**, **D**, left, **E,** left, and **F** - **I**) show confocal sections obtained at around 2-3 μm above the glass surface, whereas the confocal sections in (**D**) and (**E**), right, were captured just above the glass surface.

### Supplemental Movies

#### Supplemental Movie S1.

Mechanical stretching of a cell-sheet expressing RFP-actin observed under a real-time confocal microscope. A pair of needles was located to the left and the right, which cannot be observed in the images.

#### Supplemental Movie S2.

Confocal micrographs of a cell-sheet expressing RFP-actin before stretching. Relative distance from the first optical section is indicated in the top left corner.

#### Supplemental Movie S3.

Confocal micrographs of a cell-sheet expressing RFP-actin after stretching, held for 30 min. Relative distance from the first optical section is indicated in the top left corner.

#### Supplemental Movie S4.

Confocal micrographs of a cell-sheet expressing GFP-wt-α-catenin (green) and RFP-actin (magenta) after stretching, held for 30 min. Relative distance from the first optical section is indicated in the top left corner.

#### Supplemental Movie S5.

Confocal micrographs of a cell-sheet expressing GFP-697-α-catenin (green) and RFP-actin (magenta) after stretching, held for 30 min. Relative distance from the first optical section is indicated in the top left corner.

#### Supplemental Movie S6.

Confocal micrographs of a cell-sheet expressing GFP-β-catenin (green) and RFP-actin (magenta) after stretching, held for 30 min. Relative distance from the first optical section is indicated in the top left corner.

#### Supplemental Movie S7.

Confocal micrographs of a cell-sheet expressing GFP-vinculin (green) and RFP-actin (magenta) after stretching, held for 30 min. Relative distance from the first optical section is indicated in the top left corner.

#### Supplemental Movie S8.

Confocal micrographs of a cell-sheet expressing GFP-vinculin (green) and mCherry-β-catenin (magenta) after stretching, held for 30 min. Relative distance from the first optical section is indicated in the top left corner.

#### Supplemental Movie S9.

Confocal micrographs of a cell-sheet expressing GFP-p130Cas (green) and RFP-actin (magenta) after stretching, held for 30 min. Relative distance from the first optical section is indicated in the top left corner.

#### Supplemental Movie S10.

Confocal micrographs of a cell-sheet expressing GFP-p130Cas (green) and mCherry-β-catenin (magenta) after stretching, held for 30 min. Relative distance from the first optical section is indicated in the top left corner.

#### Supplemental Movie S11.

Confocal micrographs of a cell-sheet expressing GFP-zyxin (green) and RFP-actin (magenta) after stretching, held for 30 min. Relative distance from the first optical section is indicated in the top left corner.

#### Supplemental Movie S12.

Confocal micrographs of a cell-sheet expressing GFP-zyxin (green) and mCherry-β-catenin (magenta) after stretching, held for 30 min. Relative distance from the first optical section is indicated in the top left corner.
